# *Mycobacterium tuberculosis* Rv0991c is a redox-regulated molecular chaperone

**DOI:** 10.1101/2020.03.05.980086

**Authors:** Samuel H. Becker, Kathrin Ulrich, Avantika Dhabaria, Beatrix Ueberheide, William Beavers, Eric P. Skaar, Lakshminarayan M. Iyer, L. Aravind, Ursula Jakob, K. Heran Darwin

**Affiliations:** Department of Microbiology, New York University School of Medicine, 430 E. 29^th^ Street, Suite 312, New York, NY 10016, USA; Department of Microbiology and Immunology, University of Minnesota, 2101 6^th^ Street SE, Campus Code 2641, Minneapolis, MN 55455; Department of Molecular, Cellular, and Developmental Biology, University of Michigan, 1105 N. University Ave, Ann Arbor, MI 48109; Proteomics Laboratory, Division of Advanced Research Technologies, New York University School of Medicine, 430 E. 29^th^ Street, Suite 860, New York, NY 10016; Vanderbilt Institute of Infection, Immunology, and Inflammation, Vanderbilt University Medical Center, Nashville, Tennessee, 37037; National Center for Biotechnology Information, National Library of Medicine, National Institutes of Health, Bethesda, MD 20894, USA

## Abstract

The bacterial pathogen *Mycobacterium (M.) tuberculosis* is the leading cause of death by an infectious disease among humans. Here, we describe a previously uncharacterized *M. tuberculosis* protein, Rv0991c, as a molecular chaperone that is activated by oxidation. Rv0991c has homologues in most bacterial lineages and appears to function analogously to the well-characterized *Escherichia coli* redox-regulated chaperone Hsp33, despite a dissimilar protein sequence. Rv0991c is transcriptionally co-regulated with *hsp60* and *hsp70* chaperone genes in *M. tuberculosis*, suggesting that Rv0991c functions with these chaperones in maintaining protein quality control. Supporting this hypothesis, we found that, like oxidized Hsp33, oxidized Rv0991c prevents the aggregation of a model unfolded protein *in vitro*, and promotes its refolding by the *M. tuberculosis* Hsp70 chaperone system. Furthermore, Rv0991c interacts with DnaK and associates with many other *M. tuberculosis* proteins. Importantly, we found Rv0991c is required for the full virulence of *M. tuberculosis* in mice. We therefore propose that Rv0991c, which we named “Ruc” (redox-regulated protein with <u>u</u>nstructured <u>C</u>-terminus), represents a founding member of a new chaperone family that protects *M. tuberculosis* and other species from proteotoxicity during oxidative stress.

**IMPORTANCE:** *M. tuberculosis* infections are responsible for more than one million human deaths per year. Developing effective strategies to combat this disease requires a greater understanding of *M. tuberculosis* biology. As in all cells, protein quality control is essential for the viability of *M. tuberculosis*, which likely faces proteome stress within a host. Here, we identify an *M. tuberculosis* protein, Ruc, that gains chaperone activity upon oxidation. Ruc represents a previously unrecognized family of redox-regulated chaperones found throughout the bacterial super-kingdom. In addition to elucidating the activity of this chaperone, we found that Ruc was required for full *M. tuberculosis* virulence in mice. This work contributes to a growing appreciation that oxidative stress may provide a particular strain on protein stability in cells, and may likewise play a role in *M. tuberculosis* pathogenesis.

## INTRODUCTION

The folding of a protein that is in a non-native conformation, including during translation or upon stress-induced denaturation, can be accomplished in all organisms by a set of chaperones belonging to the Hsp70 and Hsp40 protein families. In bacteria, these chaperones are called DnaK and DnaJ, respectively [reviewed in (1)]. DnaK is an ATPase that iteratively binds to and releases protein substrates, allowing them to fold into a native conformation. In the ATP-bound state, DnaK has high substrate affinity; substrate binding subsequently induces ATP hydrolysis in a DnaJ-dependent manner. The low-affinity, ADP-bound DnaK releases the client protein, which has either reached its native structure or can re-bind to DnaK (2, 3). This cycle of high- and low-affinity substrate-binding states is facilitated by a nucleotide exchange factor (NEF), which places the DnaK nucleotide-binding cleft in an open conformation; in bacteria, this activity is accomplished by a single DnaK-associated NEF, GrpE (4). Besides harboring intrinsic protein folding activity, the DnaK/DnaJ/GrpE system (DnaKJE) can also deliver substrates to GroEL/GroES chaperonins (also known as Hsp60/Hsp10), and eukaryotic DnaKJE homologs can cooperate with proteases to promote the degradation of unfolded substrates (5–7).

Proteins in non-native conformations often expose hydrophobic regions that are prone to aggregation, an event that is toxic to cells. Importantly, aggregate formation becomes irreversible if DnaKJE cannot access unfolded proteins for refolding. To remedy this situation, the Hsp100 family of ATP-dependent chaperones (called ClpB in bacteria) cooperate with DnaKJE by solubilizing proteins within aggregates (8–10). The prevention of irreversible protein aggregation is also accomplished in all organisms by the small heat shock protein (sHsp) family of chaperones. Rather than actively dissociating protein aggregates by ATP hydrolysis, sHsps act by simply binding to denatured or misfolded proteins; the presence of sHsps within protein aggregates allows for efficient refolding by ATPase chaperones (5, 11–14). Thus, in contrast to “refoldases”, sHsps are “holdases” that afford cells the ability to rapidly respond to proteotoxic stress in an energy-independent manner.

While DnaKJE, chaperonins, ClpB, and sHsps are all active under steady-state conditions, there are also non-canonical chaperones that only become active upon encountering specific stresses. Hsp33 (encoded by *hslO*), found in both prokaryotes and eukaryotes, and the eukaryotic Get3 are inactive as chaperones in the normal reducing environment of the cytoplasm; however, oxidation of cysteine thiols within Hsp33 or Get3 induces conformational changes that allow them to bind to unfolded proteins (6, 15–18). Hsp33 prevents the irreversible aggregation of unfolded proteins by delivering substrates to ATPase chaperones for refolding (6). In *E. coli*, this activity is of particular importance during heat and oxidative stress, during which low ATP levels cause DnaK to enter a partially unfolded, nucleotide-depleted state; the presence of Hsp33 allows for refolding by DnaKJE to proceed once a normal temperature and reducing environment is restored (19). Another bacterial chaperone, RidA, is activated by the oxidant hypochlorous acid through a mechanism that does not involve oxidation of cysteine thiols, but rather chlorination of its free amino groups (20). Aside from oxidation, other non-canonical chaperones have been found to activate upon exposure to high temperatures or acid [reviewed in (21)].

The bacterial pathogen *M. tuberculosis* is currently responsible for the majority of human infectious disease-related deaths worldwide (22). In this work, we describe our discovery that the uncharacterized *M. tuberculosis* gene Rv0991c encodes a chaperone, which we named Ruc (redox-regulated chaperone with <u>u</u>nstructured <u>C</u>-terminus). Ruc belongs to a previously unacknowledged, but evolutionarily widespread family of bacterial proteins with little predicted structural similarity to other chaperones. Upon oxidation, Ruc is capable of inhibiting protein aggregation and delivering unfolded proteins to DnaKJE for refolding. Ruc was also found to interact with DnaK in *M. tuberculosis* and, while dispensable for *in vitro* growth, was required for full *M. tuberculosis* virulence in mice. These observations ultimately suggest that Ruc is important for the ability of *M. tuberculosis*, and potentially many other bacterial species, to withstand oxidation-associated proteotoxicity.

## RESULTS

### Ruc induction during heat stress requires the *M. tuberculosis* Pup-proteasome system

We became interested in Ruc after identifying the transcriptional repressor HrcA as a putative target of the *M. tuberculosis* Pup-proteasome system (23). In *M. tuberculosis*, HrcA directly represses four genes; three of them encode highly conserved chaperone proteins of the Hsp60/Hsp10 family, while the fourth gene, *ruc*, encodes a protein of unknown function (24). In addition to its negative regulation by HrcA, *ruc* is induced during heat stress by SigH, a sigma factor that also activates transcription of the Hsp70/Hsp40 genes (25). The DNA sequences to which SigH and HrcA bind upstream of the *ruc* start codon overlap, suggesting that these two transcriptional regulators compete for binding to the *ruc* promoter (**Fig. 1A**). We compared Ruc abundance upon incubation of *M. tuberculosis* at 37°C or 45°C and found that Ruc levels increased dramatically at 45°C, consistent with previously reported transcriptional data (**Fig. 1B, lanes 1 and 2**) (24). The low abundance of Ruc at 37°C is primarily due to repression by HrcA, given that Ruc was highly abundant in an *hrcA* mutant at this temperature (**Fig. 1B, lane 5**). We observed a further induction of Ruc levels during heat shock in an *hrcA* mutant (**Fig. 1B, lane 6**). These results corroborate earlier evidence that *ruc* expression is controlled in two ways: through repression by HrcA and induction by SigH.

**Fig. 1.**
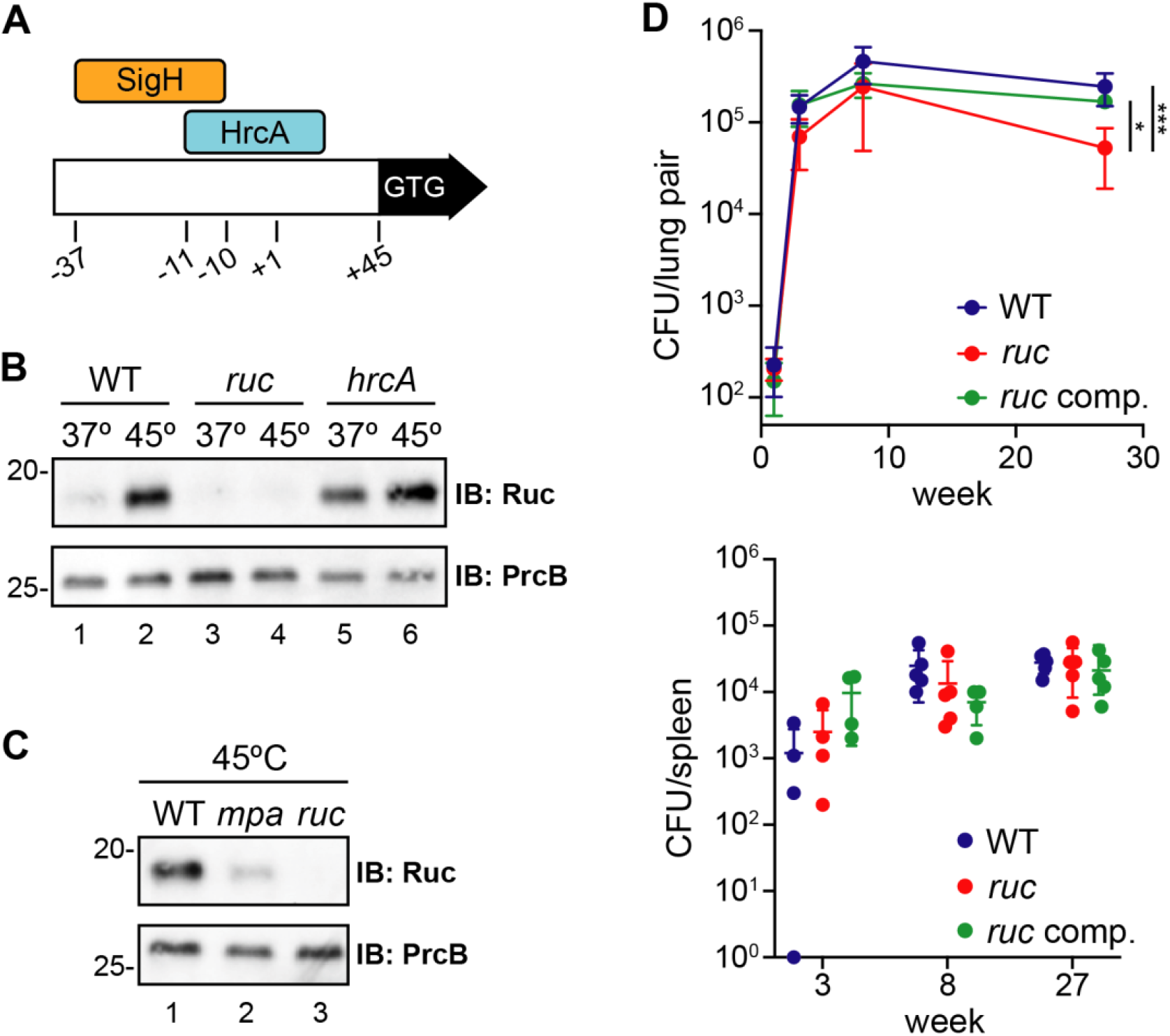
*M. tuberculosis* Ruc is a heat shock-inducible, HrcA- and Mpa-regulated small protein. **(A)** Illustration of the *ruc* control region in *M. tuberculosis*. The position of the *ruc* transcriptional start site (+1), as well as the binding sites of sigma factor SigH and repressor HrcA are shown relative to the +1 (24, 25). A second +1 was identified 19 nucleotides upstream of the +1 shown here (67). **(B)** WT (MHD1), *ruc* (MHD1384), and *hrcA* (MHD1384) *M. tuberculosis* strains were incubated at 37°C or 45°C, and Ruc abundance was assessed in bacterial lysates by immunoblot (IB). Immunoblotting for PrcB was used as a loading control. **(C)** WT, *mpa* (MHD149), and *ruc* strains analyzed as in panel (B). **(D)** Mice were infected with WT with empty vector (MHD1385), *ruc* with empty vector (MHD1393), or *ruc* complemented with pMV306kan-*ruc* (MHD1394) *M. tuberculosis* strains, and bacterial burden in the lungs (top panel) and spleen (bottom panel) was determined at day 1 or at weeks 3, 8, and 27. Statistical significance was determined using one-way ANOVA; ***, *p* < 0.001; *, *p* < 0.05. Significant differences in lung bacterial burden are shown for 27 weeks after infection; all other time points were determined to be not significant (*p* > 0.05).

In a recent study, we proposed that the pupylation and degradation of HrcA by the *M. tuberculosis* proteasome is required for the full expression of the HrcA regulon. We previously reported the abundance of HrcA-regulated gene products between a wild type (WT) *M. tuberculosis* strain and an *mpa* mutant, which cannot degrade pupylated proteins, and found that GroES, GroEL1, and GroEL2 levels are significantly lower in an *mpa* strain; however, Ruc abundance is unaffected by disruption of *mpa* under these conditions. This result suggested that proteasomal degradation of HrcA is not sufficient for Ruc production under the conditions tested (23); however, this experiment was performed with cultures incubated at 37°C in minimal medium. We therefore compared Ruc abundance from the same strains incubated at 45°C and observed a striking defect in Ruc production in the *mpa* mutant (**Fig. 1C, lane 2**). This result supports the hypothesis that the proteasomal degradation of HrcA is required for *M. tuberculosis* to robustly induce *ruc* expression during heat stress.

### Ruc is required for the full virulence of *M. tuberculosis*

Previous studies have shown that *M. tuberculosis* protein quality control pathways are important for its pathogenesis. *M. tuberculosis* strains deficient in *clpB* or the sHsp-encoding *acr2* have impaired virulence in mice (26, 27), while a mutant lacking HspR, which represses the expression of *clpB*, *acr2*, and the *hsp70/hsp40* genes, also produces less severe infections (28). Because *ruc* is co-regulated with essential chaperones, we tested if Ruc were required for the full virulence of *M. tuberculosis*. We inoculated mice with *M. tuberculosis* strains by aerosol, and assessed bacterial burden in the lungs and spleen at several time points following infection. At three- or eight-weeks post-infection, a *ruc* mutant did not display a significant survival defect compared to a WT strain. However, after 27 weeks, we recovered approximately five-fold fewer bacteria from the lungs of mice infected with the *ruc* mutant than from those infected with WT or *ruc* complemented strains (**Figure 1D, top panel**). Bacterial burden in the spleen was not significantly different between infections at any time point (**Figure 1D, bottom panel**). While the virulence phenotype of the *ruc* mutant was modest, our results indicate that Ruc may be important for *M. tuberculosis* survival in the lungs during the chronic stage of an infection.

### Ruc is part of a novel protein family found across the bacterial super-kingdom

According to the mycobacterial genome database MycoBrowser, Ruc is conserved in both pathogenic and non-pathogenic mycobacteria (29). To determine the broader phyletic distribution of Ruc among bacteria, we searched the Ruc protein sequence against a curated collection of 7,423 complete prokaryotic genomes. Ruc is highly conserved throughout the planctomycetes, chloroflexi, delta- and beta-proteobacteria, actinobacteria, spirochaetes, and verrucomicrobiae lineages; additionally, Ruc is found in species within other major bacterial lineages such as gammaproteobacteria, firmicutes and cyanobacteria. This phyletic pattern suggests that Ruc was present in the last bacterial common ancestor, with rare lateral transfers to archaea (**Table 2**).

To define the conserved features of Ruc, we used Phyre2, a program that predicts secondary and tertiary protein structures based on published sequences and solved structures (30). Alignment of Ruc with its homologs in other bacteria identified two conserved domains. The amino (N)-terminal region, consisting of approximately 50 amino acids, contains four cysteine (Cys) residues arranged in a manner consistent with a zinc ribbon fold, a domain found in zinc-binding proteins of highly diverse functions (**Fig. 2A**) (31). The carboxyl (C)-terminal regions of Ruc homologs consist of highly variable sequences of amino acids ranging in length from approximately seven to 61 residues. While these C-terminal domains do not share significant similarity at the protein sequence level, they appear to be uniformly composed of hydrophilic residues with no predicted secondary structure. Overall, these structural predictions allowed us to conclude that *M. tuberculosis* Ruc and its homologs are likely characterized by a metal-coordinating N-terminal domain and a disordered C-terminus (**Fig. 2B**).

**Fig. 2.**
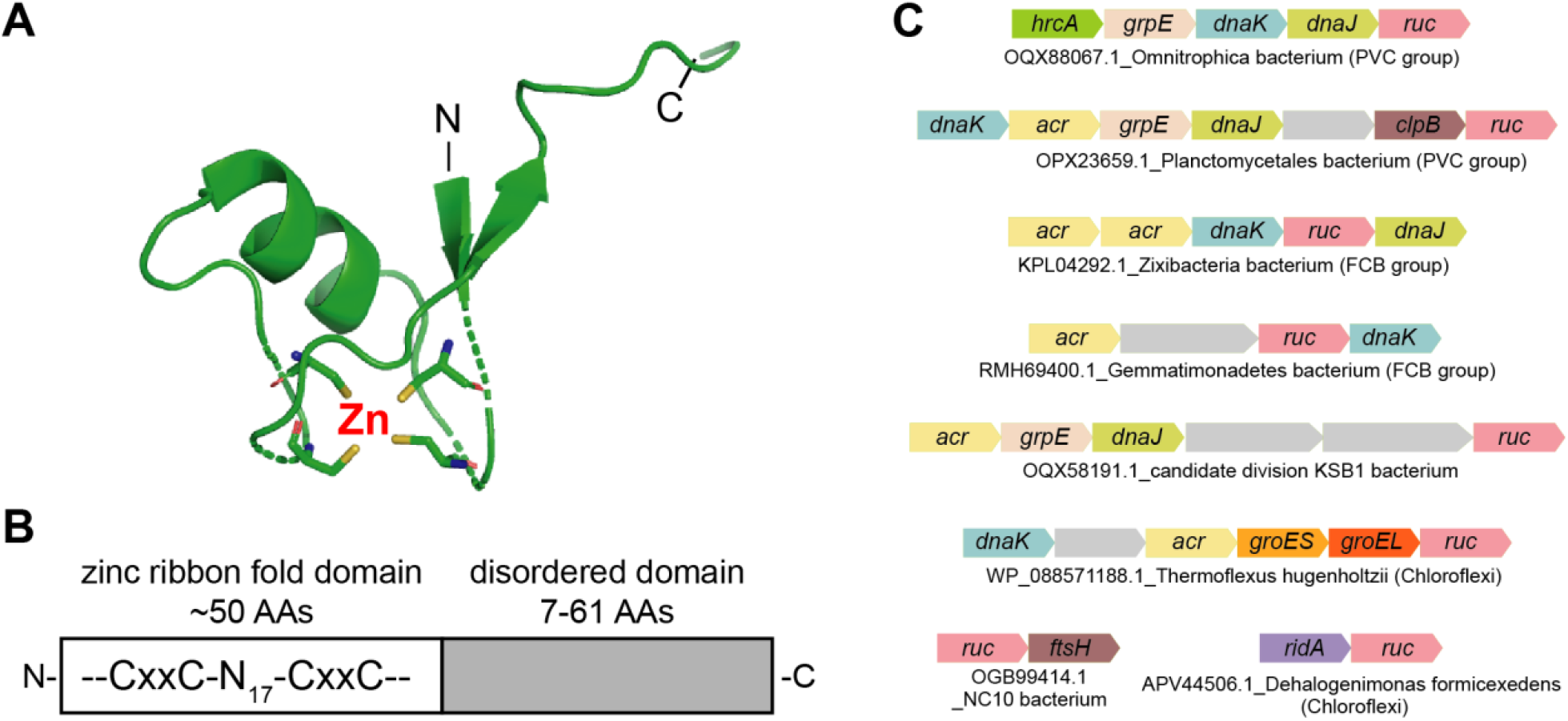
Ruc contains a putative zinc-binding domain with a disordered C-terminus, and co-occurs with proteostasis genes in diverse bacterial lineages. **(A)** Predicted structure of the N-terminal region of *M. tuberculosis* Ruc (residues 5-48) based on Phyre2 predictive modeling (30). The four cysteines in Ruc are represented as sticks, with the thiol groups shown in yellow. The position of a predicted zinc ion is also shown. **(B)** Illustration of the conserved features of Ruc. x, unspecified amino acid; N_17_, region 17 residues in length. **(C)** Genetic loci containing *ruc* with neighboring genes encoding chaperones, proteases, or chaperone-associated transcriptional regulators. Representatives of the phyletic groups PVC (Plantomycetes, Verrucomicrobia, Chlamydiae) and FCB (Fibrobacteres, Chlorobi, Bacteroidetes), as well as members of Chloroflexi and unclassified phyla, are shown. Genes are represented by the NCBI GenBank database accession number of the *ruc* gene followed by the species name and bacterial clade in bracket (if known).

Having established that *ruc* is co-expressed with the Hsp60 and Hsp70 chaperone system genes in *M. tuberculosis* and is present in diverse bacterial lineages, we next asked if *ruc* is associated with protein quality control genes in other species. A comprehensive gene neighborhood analysis among bacteria recovered widespread associations with chaperone genes, as well as transcription factors and proteases that regulate chaperone production. In bacteria of the PVC (Plantomycetes, Verrucomicrobia, Chlamydiae) and FCB (Fibrobacteres, Chlorobi, Bacteroidetes) superphyla, *ruc* is present in loci encoding DnaK, DnaJ, GrpE, ClpB, alpha-crystallin sHsps (Acr), and HrcA. In bacteria from the Chloroflexi phylum, as well as several unclassified species, Ruc is additionally associated with GroES/GroEL chaperonins, the chaperone-regulating protease FtsH, and the redox-regulated chaperone RidA (**Fig. 2C**). The identification of these conserved genetic linkages, as well as the observation that *ruc* is transcriptionally co-regulated with essential chaperone genes in *M. tuberculosis*, allowed us to conclude that Ruc most likely performs a function related to protein folding.

### Ruc coordinates zinc and is an intrinsically disordered protein

Based on the observations that *ruc* is closely associated with essential chaperone genes and that Ruc contains putative zinc-coordinating cysteines, we hypothesized that Ruc has oxidation-dependent chaperone activity. Hsp33 and Get3, the only chaperones currently described whose activity requires cysteine oxidation, each contain a zinc-coordinating domain consisting of four cysteines. Under oxidizing conditions, these cysteines form intramolecular disulfide bonds and release zinc, allowing these chaperones to form complexes with denatured proteins to prevent their irreversible aggregation (15, 17, 32).

To begin to test Ruc chaperone activity using *in vitro* assays, we produced and purified *M. tuberculosis* Ruc from *E. coli* under reducing conditions (Ruc_red_). We oxidized purified Ruc (Ruc_ox_) by incubation with hydrogen peroxide (H_2_O_2_) and copper chloride (CuCl_2_), which react to generate hydroxyl radicals that rapidly oxidize cysteines (33). On an SDS-PAGE gel, Ruc_red_ migrated as a single band (**Fig. 3A, lane 1**) while Ruc_ox_ migrated through the gel as multiple species, whose sizes were consistent with the formation of covalent, intermolecular disulfide bonds (**Fig. 3A, lane 2**). Treatment of Ruc_ox_ with the thiol-reducing agent dithiothreitol (DTT) resulted in a significant loss of higher-molecular weight species, indicating that the oxidation of Ruc cysteines is reversible (**Fig. 3A, lane 3**). These results demonstrate that Ruc cysteines form disulfide bonds under oxidizing conditions to create covalently-linked multimers.

**Fig. 3.**
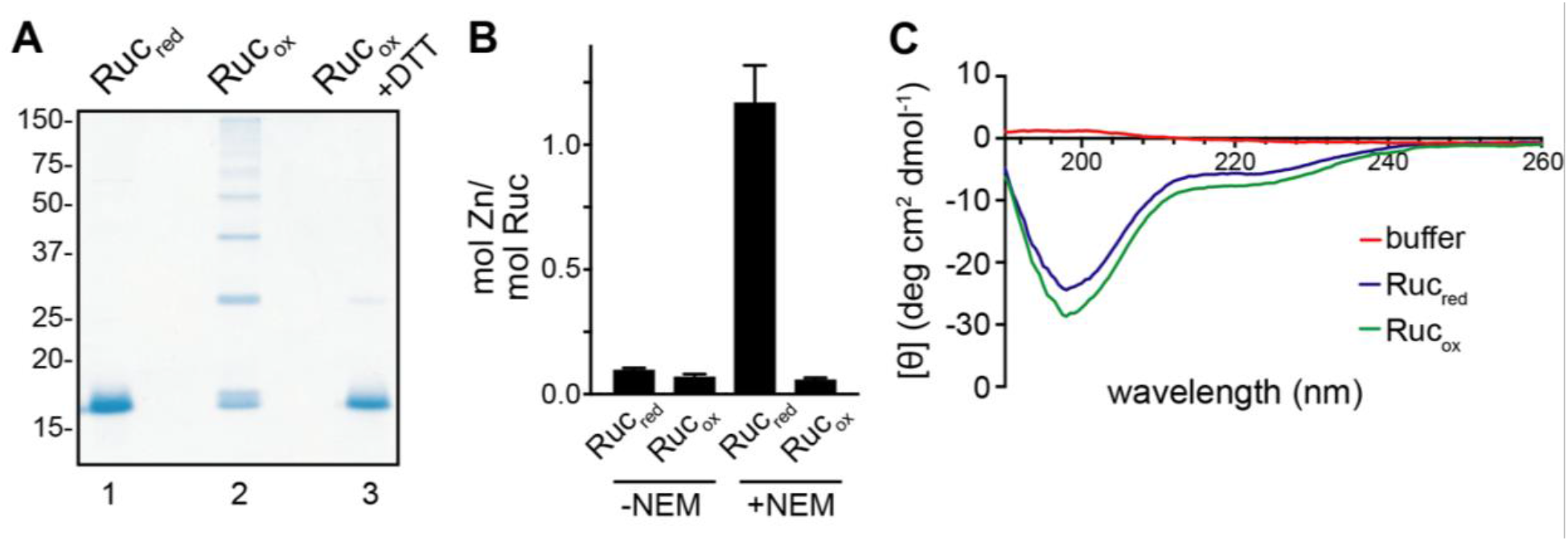
Ruc contains redox-active cysteines that coordinate a single zinc atom, and is an intrinsically disordered protein. **(A)** Reduced Ruc (Ruc_red_), oxidized Ruc (Ruc_ox_), or Ruc_ox_ treated with the thiol-reducing agent dithiothrietol (DTT) were separated on an SDS-PAGE gel and stained with Coomassie brilliant blue. **(B)** Zinc coordination by Ruc_red_ or Ruc_ox_ was quantified in the absence or presence of N-ethylmaleimide (NEM), which modifies cysteines. Zinc concentrations were measured spectrophotometrically using the metal chelator 4-(2-pyridylazo)resorcinol (PAR) (see Materials and Methods for details). **(C)** Assessment of Ruc secondary structure using circular dichroism, with degrees of ellipticity [θ] plotted by wavelength.

In Hsp33, four conserved cysteines coordinate zinc. This zinc binding maintains Hsp33 in an inactive state, yet allows Hsp33 to become readily activated once an oxidant is present (32). To determine if Ruc binds zinc using its four cysteines, we used 4-(2-pyridylazo)resorcinol (PAR), a chemical that chelates free zinc to yield an absorbance peak at 500 nm (A_500_) (see Materials and Methods). Incubation of either Ruc_red_ or Ruc_ox_ with PAR alone did not yield a significant change in A_500_, demonstrating that the protein preparations contained little free zinc; however, addition of N-ethylmaleimide (NEM), which forms adducts on cysteine thiols, resulted in the release of zinc from Ruc_red_ that could be detected in approximately equimolar abundance to Ruc_red_. Meanwhile, addition of NEM to Ruc_ox_ did not reveal any change in zinc levels (**Fig. 3B**). These data suggest that under reducing conditions, Ruc cysteines coordinate an atom of zinc, and that this binding is disrupted when the cysteine thiols are oxidized.

Given that PAR can chelate metals in addition to zinc (34), we next determined the precise identity of the metal bound to Ruc using inductively coupled plasma mass spectrometry (ICP-MS). In this analysis, we found zinc was present in Ruc_red_ preparations in close to equimolar amounts; when Ruc_red_ was treated with NEM prior to ICP-MS, little Ruc-bound zinc was detected (**Table S1**). Taken together, our metal-binding assays demonstrated that cysteine thiols in Ruc coordinate a single zinc atom.

Aside from the N-terminal zinc ribbon fold domain, the entire C-terminal half of Ruc was predicted to lack secondary structure (**Fig. 2B**). To evaluate the degree of disorder in Ruc, we measured the secondary structures found in Ruc_red_ and Ruc_ox_ using circular dichroism (CD) spectroscopy [reviewed in (35)]. In accordance with structural predictions, both Ruc_red_ and Ruc_ox_ yielded a CD spectrum characteristic of disordered proteins (**Fig. 3C**) (36). The high degree of disorder present in Ruc, along with our previous observation that all Ruc homologs are predicted to harbor domains that lack secondary structure, support a hypothesis whereby the unstructured C-terminal domain somehow contributes to the function of this protein.

### Oxidized Ruc prevents unfolded protein aggregation *in vitro*

Chaperones are able to inhibit protein aggregation due to their propensity to bind to unfolded proteins. A common method for detecting chaperone activity uses purified firefly luciferase, which denatures and forms irreversible aggregates when heated above 42°C; the presence of a chaperone during denaturation of luciferase prevents its aggregation (15, 37). We therefore used this method to test chaperone activity by Ruc. When we incubated luciferase at 45°C, we observed the formation of precipitates that could be detected by an increase in light absorbance at 350 nm. In the presence of a fivefold molar excess of Ruc_red_, we observed a similar level of luciferase aggregation, demonstrating that Ruc_red_ has little to no chaperone activity. In contrast, the presence of Ruc_ox_ during heat inactivation, either in excess or at an equimolar concentration, significantly inhibited luciferase aggregation (**Fig. 4A**). When we measured luciferase aggregation using a different method of detection, dynamic light scattering (38), we also observed the inhibition of aggregation by Ruc_ox_, but not by Ruc_red_ (**Fig. S1A**). Furthermore, chaperone activity by Ruc was observed when Ruc was pre-treated with the oxidizing agents hypochlorite or nitric oxide (**Fig. S1B**), further supporting a model whereby oxidized Ruc counteracts protein aggregation.

**Fig. 4.**
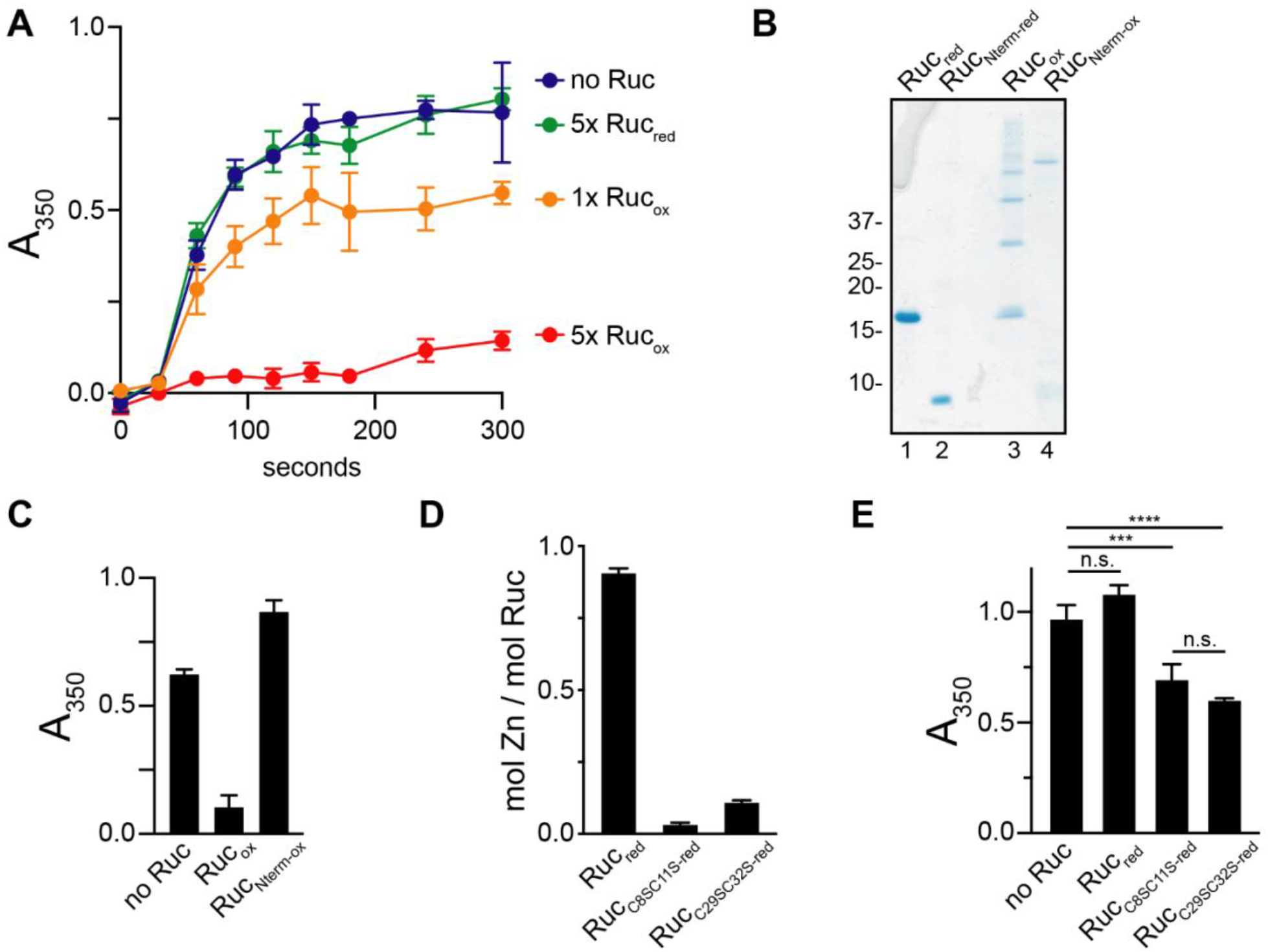
Oxidized Ruc inhibits protein aggregation. **(A)** Aggregation of luciferase upon heat denaturation. Luciferase was incubated at 45°C either alone or in the presence of a five-fold molar excess (5x) of Ruc_red_, 5x Ruc_ox_, or an equimolar concentration (1x) of Ruc_ox_. Aggregation was assessed by absorbance at 350 nm (A_350_). The difference in aggregation between no Ruc and 1x Ruc_ox_ or 5x Ruc_ox_ conditions was statistically significant (paired *t*-test, *P* < 0.01), while no significant difference was obtained with Ruc_red_. **(B)** Ruc and Ruc_Nterm_ (comprising residues 1-49), in either a reduced or oxidized state, were separated on a Coomassie-stained SDS-PAGE gel. **(C)** Aggregation of heat-denatured luciferase as in (A), except only the 300 second time point is shown. Native Ruc_ox_ or Ruc_Nterm-ox_ were incubated with luciferase in fivefold molar excess at 45°C as indicated. **(D)** Quantification of zinc in native Ruc and Ruc cysteine-to-serine variants, as described for Figure 3B. NEM was included in all reactions. **(E)** Luciferase aggregation assay as in (C) to assess the activity of reduced Ruc cysteine-to-serine variants. Statistical significance was determined using one-way ANOVA; ****, *P* < 0.0001; ***, *P* < 0.001; n.s., not statistically significant (*P* > 0.05). All reactions were performed in triplicate.

In the activation process of Hsp33, oxidation induces conformational changes that expose a disordered region with high affinity for client proteins (39, 40). We therefore asked if the intrinsically disordered C-terminal domain of Ruc was required for its chaperone activity. We produced a truncated Ruc variant, Ruc_Nterm_ (amino acids 1 through 49), which harbors only the zinc binding motif (**Fig. 4B, lane 2**). Oxidized Ruc_Nterm_ (Ruc_Nterm-ox_) formed a single high-molecular weight species, in contrast to the variety of multimers observed for Ruc_ox_ (**Fig. 4B, lanes 3 and 4**). When we tested the chaperone activity of oxidized Ruc_Nterm_ (Ruc_Nterm-ox_), we found that it was unable to prevent luciferase aggregation (**Fig. 4C**). Thus, the disordered C-terminus of Ruc is required for its chaperone activity, either by binding to client protein, by influencing the conformation or oligomerization state of Ruc, or through a combination of factors.

We next asked if replacing the Ruc cysteines, which we predicted would disrupt zinc coordination, would allow for constitutive Ruc activity in the absence of oxidation, a phenomenon that was observed for Hsp33 (32). We made cysteine (C) to serine (S) substitutions in Ruc, generating Ruc_C8S,C11S_ and Ruc_C29S,C32S_ (see Table 1 for primers and plasmids). Neither Ruc_C8S,C11S-red_ nor Ruc_C29S,C32S-red_ were bound to zinc upon their purification (**Fig. 4D**). When we tested the ability of each reduced Ruc variant to inhibit luciferase aggregation, we found that both exhibited significant chaperone activity compared to wild-type Ruc_red_ (**Fig. 4E**). Collectively, these results support a model by which Ruc is in a zinc-bound, inactive state under reducing conditions, and that oxidation and zinc release promotes a conformation of Ruc that allows for its interaction with an unfolded protein.

**Table 1.**
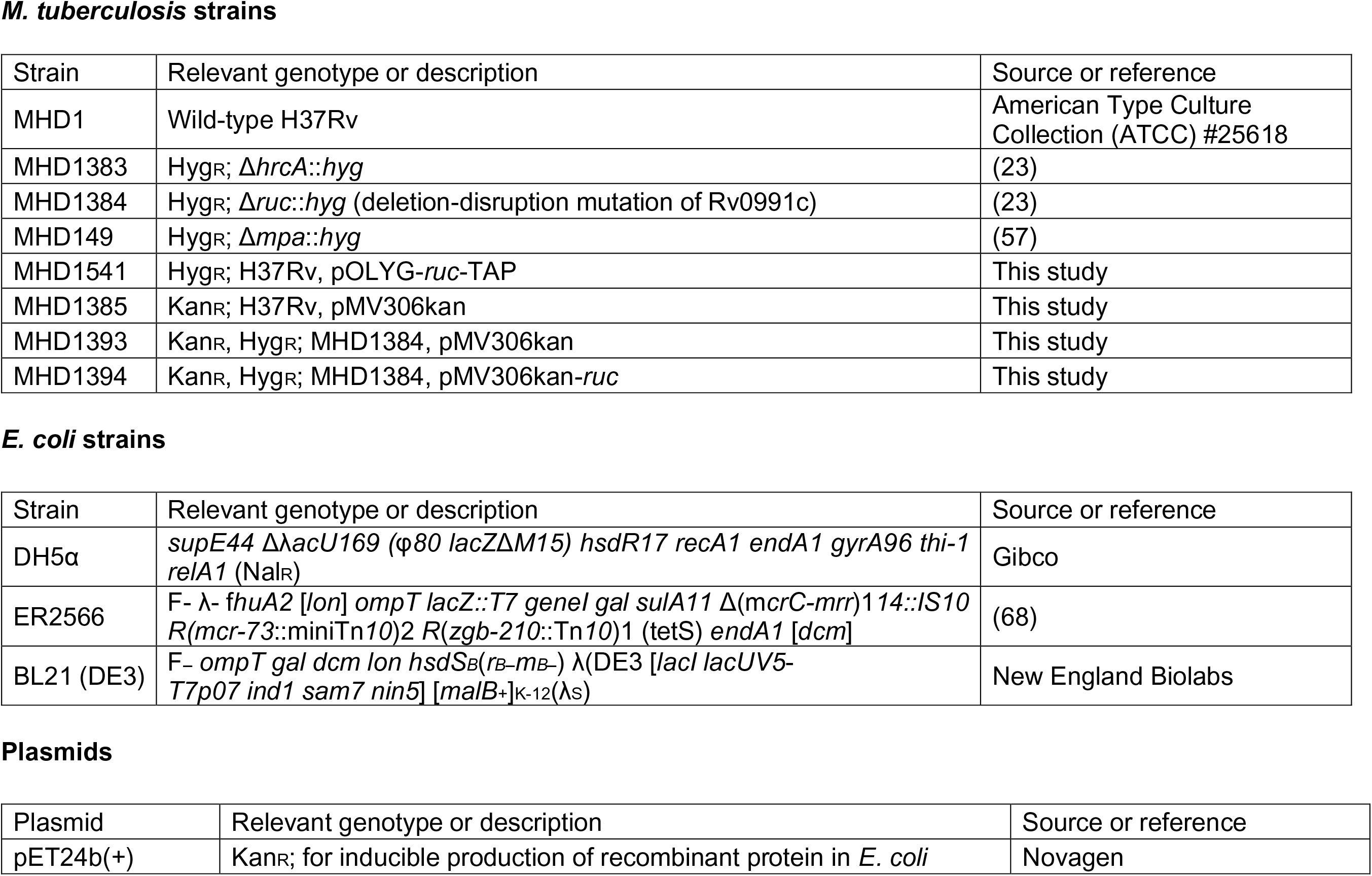
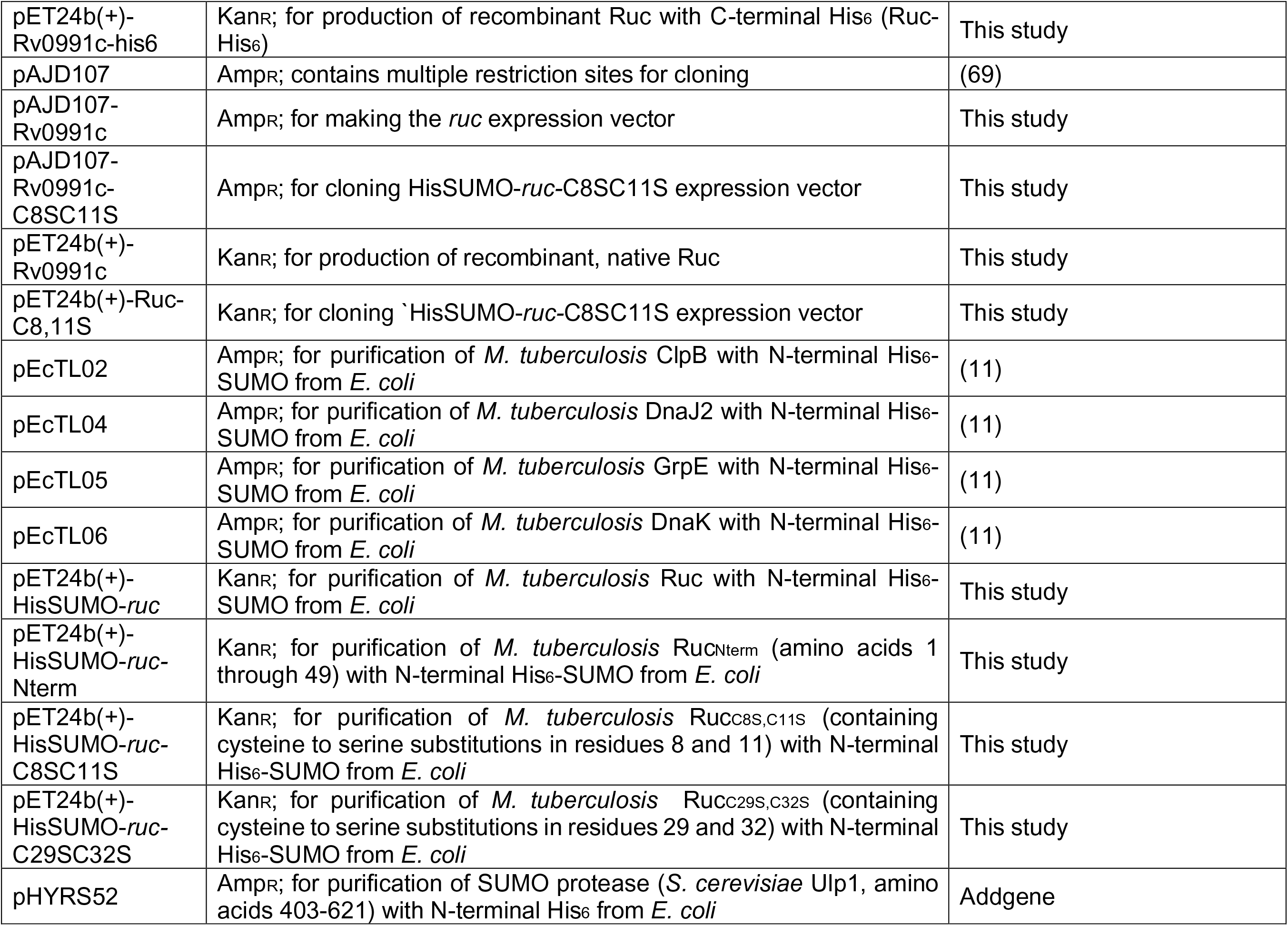
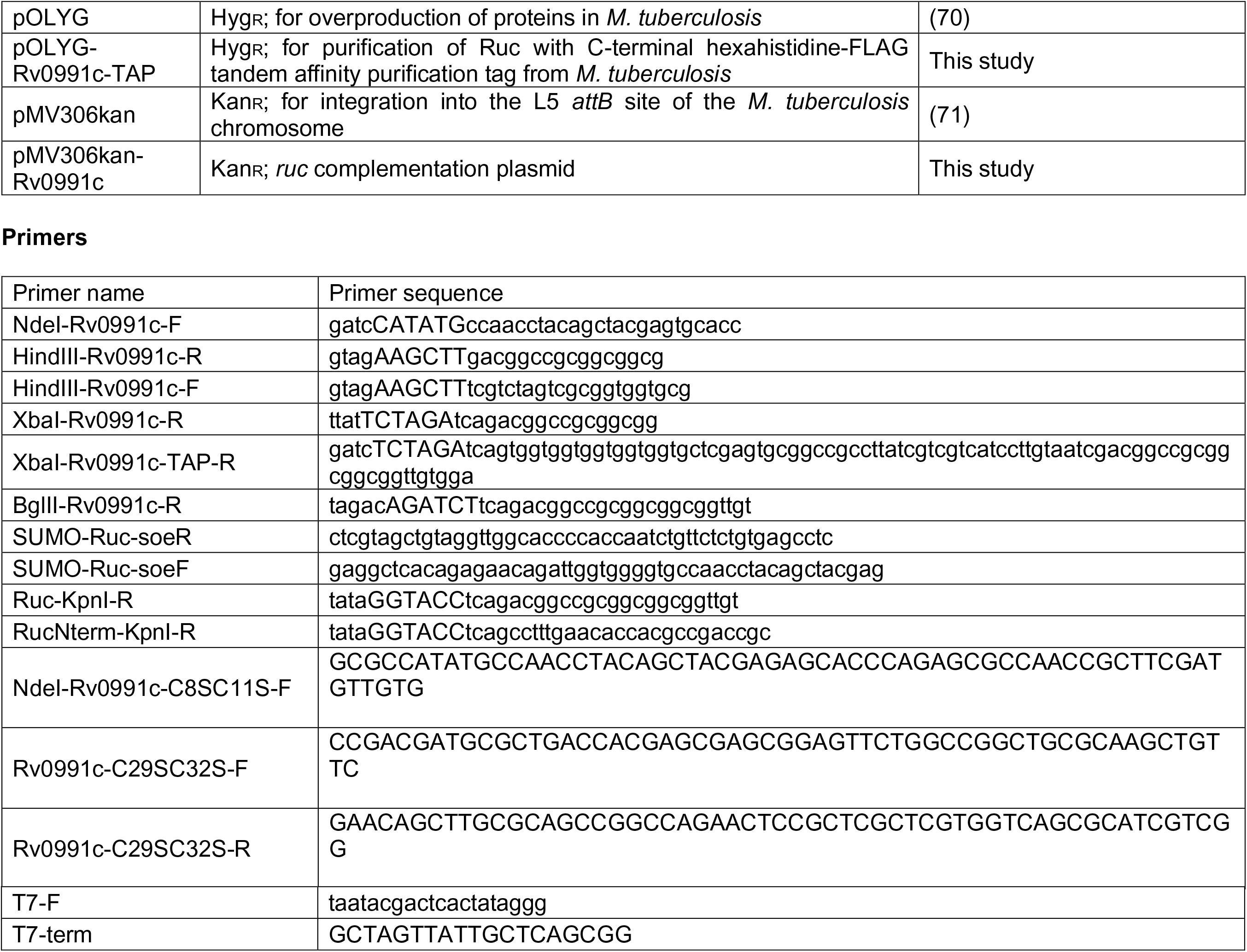
Strains, plasmids and primers used in this work.

### Oxidized Ruc promotes protein refolding by *M. tuberculosis* Hsp70/Hsp40 chaperones

Non-ATPase bacterial chaperones such as Hsp33 and sHsps prevent irreversible protein aggregation *in vivo*; critically, this function relies on the ability of ATP-dependent chaperones to eventually refold substrates that are bound by these non-ATPase chaperones (5, 6, 41). To further understand the potential role of Ruc in *M. tuberculosis* protein quality control, we tested if Ruc could promote protein refolding by *M. tuberculosis* DnaKJE. To test this hypothesis, we heated luciferase either alone or in the presence of Ruc_red_ or Ruc_ox_; after bringing the reaction to room temperature and adding purified *M. tuberculosis* DnaK, DnaJ2, and GrpE under reducing conditions, we monitored refolding by measuring luciferase activity over time. Only minimal refolding of luciferase by DnaKJE was observed when luciferase was denatured in the absence of a chaperone, a result that was expected based on previous studies (5, 6, 11, 41). By contrast, significant DnaKJE-mediated refolding of luciferase was achieved when luciferase was denatured in the presence of Ruc_ox_ (**Fig. 5A**). Consistent with its inability to prevent luciferase aggregation (**Fig. 4A**), Ruc_red_ had no significant effect on luciferase refolding (**Fig. 5A**). These results demonstrate that unfolded proteins that are bound by Ruc under oxidizing conditions can be delivered to the DnaKJE system for refolding.

**Fig. 5.**
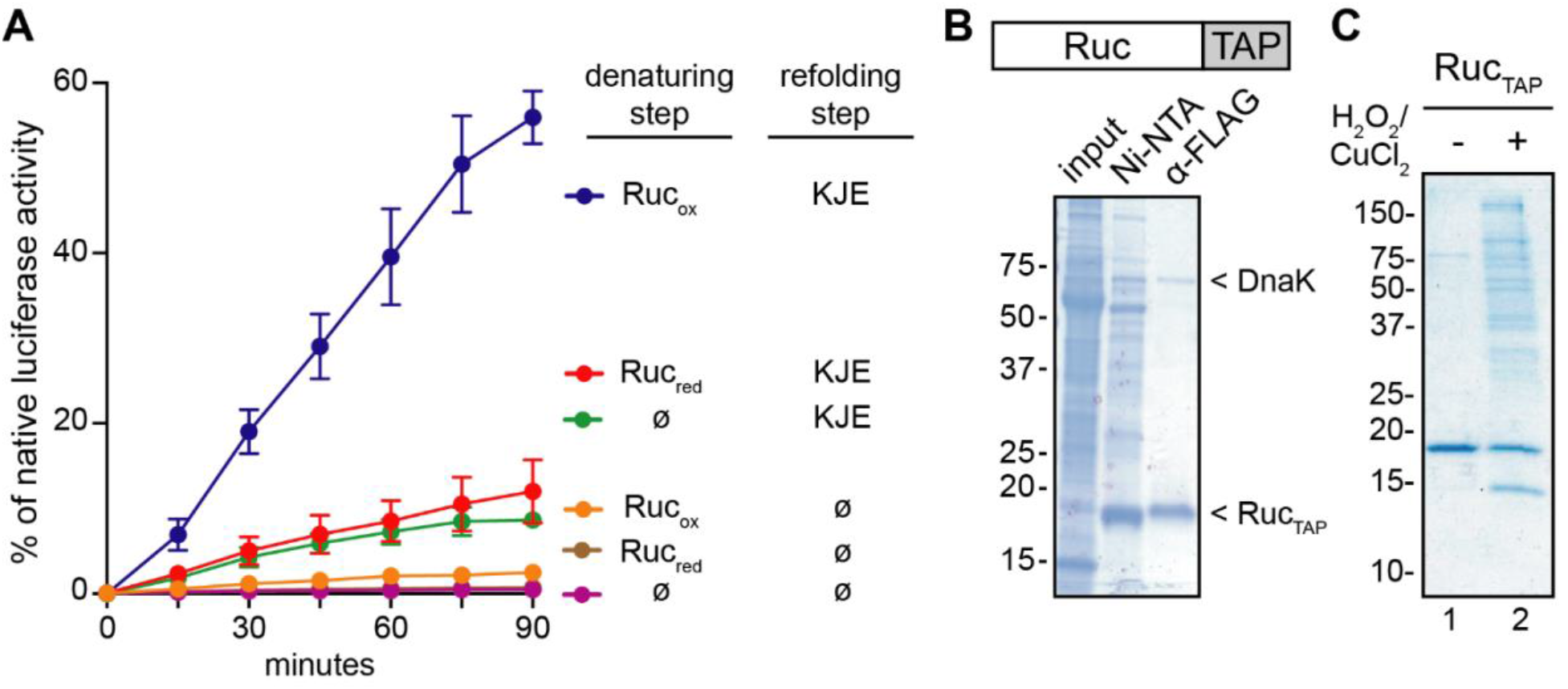
Ruc promotes protein folding by the *M. tuberculosis* Hsp70 system and associates with many *M. tuberculosis* proteins. **(A)** Luciferase was denatured at 45°C in the presence of either Ruc_red,_ Ruc_ox_, or a buffer control (ø). Reactions were then cooled to 25°C and incubated either with DnaK, DnaJ2, and GrpE (KJE). Refolding of denatured luciferase was determined by measuring luciferase activity at the indicated time points following addition of KJE or buffer. As a control, native luciferase activity was measured at each time point by incubating non-denatured luciferase in the same buffer for the same duration (see Materials and Methods for a detailed protocol). Data shown are the result of three independent experiments comparing each condition. **(B)** Ruc containing a C-terminal hexahistidine-FLAG tandem affinity purification tag (Ruc_TAP_, illustrated above) was purified from *M. tuberculosis* strain MHD1541. A two-step purification was performed on soluble *M. tuberculosis* lysates (input) using Ni-NTA resin followed by FLAG antibody gel (α-FLAG). The identity of DnaK was determined using mass spectrometry (see Materials and Methods) and immunoblotting (see Figure S2A). **(C)** Ruc_TAP_ was purified from *M. tuberculosis* as in (B), except that input lysates were subjected to oxidation (H_2_O_2_ and CuCl_2_ treatment) or no treatment prior to purification. The final α-FLAG-purified material is shown. For (B) and (C), samples were separated on SDS-PAGE gels under reducing conditions.

After establishing the chaperone activity of Ruc *in vitro*, we sought to better understand the specific role of Ruc in *M. tuberculosis* physiology by identifying potential protein interaction partners *in vivo*. To capture such interactions, we produced Ruc with a hexahistidine-FLAG tandem affinity purification tag (Ruc_TAP_) in *M. tuberculosis*. When we performed a two-step purification of Ruc_TAP_ from *M. tuberculosis* lysates under native conditions, we observed that a second protein co-purified with Ruc_TAP_ (**Fig. 5B**); mass spectrometry identified this protein as DnaK (see Materials and Methods). We validated the identity of DnaK in Ruc_TAP_ purifications by immunoblotting with a monoclonal antibody to DnaK (**Fig. S2A**). Importantly, this result was specific to Ruc_TAP_, as DnaK did not co-purify with another small TAP-tagged protein, PrcB_TAP_ (**Fig. S2A**). This apparent interaction between Ruc and DnaK in *M. tuberculosis* lysates supports the hypothesis that Ruc is directly involved in maintaining protein folding by the DnaKJE system *in vivo*.

Notably, Ruc_TAP_ interacted with DnaK under native purification conditions in which no oxidants were added. Given that Ruc chaperone function is activated under oxidizing conditions, we next sought to capture the interaction of Ruc with endogenous client proteins by purifying Ruc_TAP_ from bacteria subjected to oxidative stress, a condition that could promote association of Ruc_TAP_ with unfolded substrates. However, when we purified Ruc_TAP_ from *M. tuberculosis* cultures that were exposed to a sub-lethal concentration of H_2_O_2_ and CuCl_2_ at 45°C, we did not observe any additional proteins co-purifying with Ruc_TAP_ (**Fig. S2B**). We hypothesized that intact *M. tuberculosis* rapidly reverses oxidation, such that interactions between Ruc_ox_ and unfolded client proteins are too transient to capture. We instead treated lysates of Ruc_TAP_-producing *M. tuberculosis* with H_2_O_2_ and CuCl_2_, and found that oxidation resulted in the association of many *M. tuberculosis* proteins with Ruc_TAP_ (**Fig. 5C, lane 2**). To determine the identity of these putative clients of Ruc, we performed proteomic mass spectrometry to identify the proteins that co-purified with Ruc_TAP_ under oxidizing, but not native conditions (**Table S2**). Proteins that co-purified with activated Ruc_TAP_ were associated with a diverse range of functions including nitrogen and carbon metabolism, electron transport, oxidative stress, and transcriptional regulation; thus, Ruc chaperone activity likely provides a general protective effect on the *M. tuberculosis* proteome during oxidation. Importantly, while these results suggest that Ruc can interact with a wide variety of proteins, we are currently unable to verify that the same interactions take place within bacteria under the same conditions.

### Ruc is not essential for *M. tuberculosis* resistance to oxidation

During an infection, *M. tuberculosis* primarily resides within macrophages. These and other immune cells are capable of mounting an antimicrobial response that includes the generation of reactive oxygen species (ROS), reactive nitrogen intermediates (RNI), and hypochlorite. These molecules, all of which can react with cysteine thiols, can confer lethal stress upon infecting pathogens [reviewed in (42)]. Animals defective in the ability to produce ROS, RNI, and hypochlorite are more susceptible to bacterial and fungal pathogens (43–46), and human deficiency in ROS production is associated with susceptibility to mycobacterial infections (47).

Our data thus far led us to hypothesize that Ruc contributes to *M. tuberculosis* virulence by protecting bacteria from protein aggregation during oxidative stress. Therefore, we sought to test if a *ruc* mutant was more susceptible to a variety of oxidative stress conditions *in vitro*. We challenged *M. tuberculosis* strains incubated at 45°C with either peroxide, hypochlorite, plumbagin (which generates superoxide radicals in cells) (48), or acidified nitrite (which produces NO) (49) and measured the approximate lethal dose of each compound. Interestingly, the WT and *ruc* mutant strains were equally susceptible to killing under all stress conditions tested (**Fig. S3**). Thus, despite the observation that the *ruc* mutant has a subtle virulence defect in mice, we were unable to conclusively determine if Ruc protects *M. tuberculosis* from the various oxidative stress conditions that might be encountered in a host. Given that *M. tuberculosis* has multiple mechanisms for maintaining bacterial redox balance, including thioredoxins, mycothiol and catalase [reviewed in (50)], it is possible that the contribution of Ruc to *M. tuberculosis* fitness only becomes apparent under conditions in which these antioxidant systems are not fully effective; notably, the requirement for Hsp33 for *E. coli* resistance to oxidative stress is observed only after the thioredoxin system is genetically disrupted (16).

## DISCUSSION

In this study, we identified Ruc as the founding member of a new family of bacterial redox-regulated chaperones. Prior to this work, Hsp33 and RidA were the only other proteins described in bacteria whose chaperone activity is dependent on oxidation. Remarkably, however, Ruc shares no sequence similarity to these proteins aside from the presence of four zinc-coordinating cysteines, a feature of Hsp33. Reduced, inactive Hsp33 compactly folds into two globular domains; a combination of high temperature and oxidation causes partial unfolding of the protein, exposing a disordered region with high affinity for substrates (39, 40). The Hsp33 zinc-binding motif, while essential for inducing conformational changes upon oxidation, does not directly participate in substrate binding (39). In contrast to Hsp33, Ruc likely has a single, small globular region, and is predicted to be intrinsically disordered across more than half the length of the protein. We therefore expect that the mechanism of its activation is distinct from that of Hsp33. Our observation that Ruc_red_ has no chaperone activity could suggest that the disordered domain of Ruc_red_ is kept in a partially occluded state, perhaps by interacting with other regions of the protein, and would therefore be unavailable for binding a client protein until oxidation takes place. Structural studies of Ruc, both alone and in complex with a substrate, will be necessary to determine the precise mechanism of its activation.

It is well-established that Hsp33 and sHsps deliver unfolded proteins to ATPase chaperones for refolding (5, 6, 41). Here, we describe a similar function of Ruc in promoting protein folding by *M. tuberculosis* DnaKJE (**Fig. 6**). Importantly, despite strong evidence that proteins can be directly transferred from non-ATPase to ATPase chaperones, a direct interaction between a holdase and a foldase has never been observed (14). Thus, our ability to capture the interaction of Ruc and DnaK, even in the absence of an environmental stress, may present a new opportunity to understand mechanisms by which holdases transfer their substrates to DnaK.

**Fig. 6.**
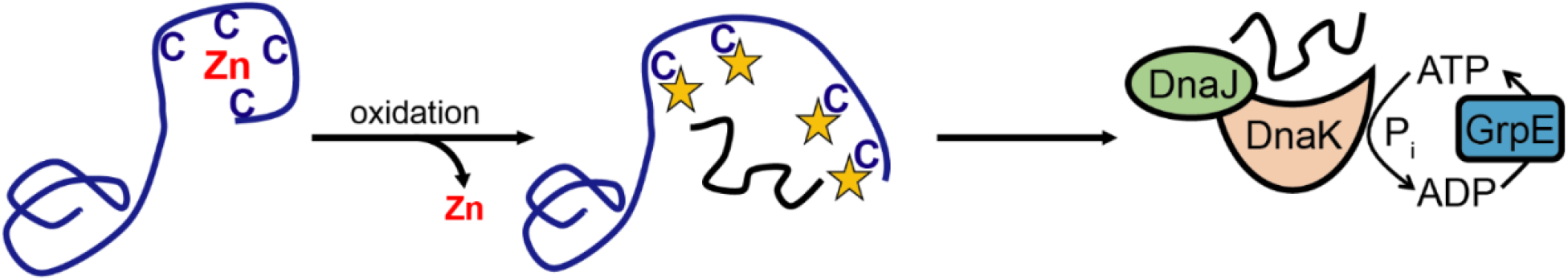
Model of Ruc chaperone activity in *M. tuberculosis*. From left to right; under the steady-state reducing conditions of the cytoplasm, Ruc coordinates zinc and is inactive; upon oxidation of Ruc cysteines, zinc is displaced and Ruc binds to unfolded proteins; a Ruc client protein is transferred to the Hsp70 system for refolding into its native conformation.

Supporting the hypothesis that Ruc contributes to *M. tuberculosis* pathogenesis, a previous screen to identify genes required for *M. tuberculosis* virulence in mice found that bacteria with transposon insertions in *ruc* are defective for *in vivo* growth during mixed infections (51). In another study, mice infected with an *M. tuberculosis* strain containing a deletion-disruption of Rv0990c (encoding Hsp22.5), the gene directly downstream of *ruc*, had a lower bacterial burden during the later stages of infection and increased time to death compared to mice infected with a WT strain; however, the phenotypes in this study could not be fully complemented (52). Conceivably, the Rv0990c mutation could have affected Ruc production, which might explain the incomplete complementation of the Rv0990c mutation to fully restore virulence.

The widespread conservation of Ruc homologues suggests that this protein protects bacteria against one or more common environmental stresses, and not just host-exclusive factors. Notably, with the exception of a single species, Hsp33 homologs are absent from the Actinobacteria; by contrast, Ruc homologs are found throughout this phylum, including the majority of mycobacterial species [STRING database (53)]. We therefore speculate that for these species, Ruc fulfills a similar role to that of Hsp33: preventing the irreversible aggregation of unfolded proteins during oxidative stress, a condition in which ATP depletion or direct thiol modification of DnaK may render DnaKJE-mediated refolding impossible (6, 19, 54). If this scenario were indeed supported by further studies, then Ruc and Hsp33, which appear to be structurally unrelated, may represent products of convergent evolution.

## Materials and Methods

### Strains, plasmids, primers, and culture conditions

See Table 1 for strains, plasmids, and primers used in this work. Reagents used for making all buffers and bacterial media were purchased from Thermo Fisher Scientific, unless otherwise indicated. *M. tuberculosis* was grown in “7H9c” [BD Difco Middlebrook 7H9 broth with 0.2% glycerol and supplemented with 0.5% bovine serum albumin (BSA), 0.2% dextrose, 0.085% sodium chloride, and 0.05% Tween-80]. For the experiment in Figure 1C, bacteria were grown in Proskauer-Beck minimal medium supplemented with asparagine and a similar result was observed in 7H9c. For solid media, *M. tuberculosis* was grown on “7H11” agar (BD Difco Middlebrook 7H11) containing 0.5% glycerol and supplemented with 10% final volume of BBL Middlebrook OADC Enrichment. For selection of *M. tuberculosis*, the following antibiotics were used as needed: kanamycin 50 µg/ml, hygromycin 50 µg/ml. *E. coli* was cultured in BD Difco Luria-Bertani (LB) broth or on LB Agar. Media were supplemented with the following antibiotics as needed: kanamycin 100 µg/ml, hygromycin 150 µg/ml, ampicillin 200 µg/ml.

Primers used for PCR amplification or sequencing were purchased from Life Technologies and are listed in Table 1. DNA was PCR-amplified using either Phusion (New England Biolabs; NEB), Pfu (Agilent), or Taq (Qiagen) according to the manufacturers’ instructions. PCR products were purified using the QIAquick Gel Extraction Kit (Qiagen). Plasmids encoding His_6_-SUMO-Ruc, His_6_-SUMO-Ruc_Nterm_, His_6_-SUMO-Ruc_C8S,C11S_ and His_6_-SUMO-Ruc_C29S,C32S_ were made using splicing by overlap extension (SOE) PCR (55). Restriction enzymes and T4 DNA ligase were purchased from NEB. The following plasmids were made by PCR-amplifying genes from *M. tuberculosis* DNA using the indicated primers and cloning amplification products into their respective vectors: pET24b(+)-Rv0991c-his_6_ (NdeI-Rv0991c-F, HindIII-Rv0991c-R), pMV306kan-Rv0991c (HindIII-Rv0991c-F, XbaI-Rv0991c-R), pOLYG-Rv0991c-TAP (HindIII-Rv0991c-F, XbaI-Rv0991c-TAP-R), pAJD107-Rv0991c (NdeI-Rv0991c-F, BglII-Rv0991c-R), pAJD107-Rv0991c-C8S,C11S (NdeI-Rv0991c-C8S,C11S-F, BglII-Rv0991c-R). pET24b(+)-Rv0991c and pET24b(+)-Rv0991c-C8S,C11S were made by subcloning the Rv0991c gene from pAJD107-Rv0991c and pAJD107-Rv0991c-C8S,C11S into pET24b(+), respectively. pET24b(+)-HisSUMO-*ruc* was made by first PCR-amplifying HisSUMO from pEcTL02 using primers T7-F and SUMO-Ruc-soeR, secondly amplifying *ruc* from *M. tuberculosis* DNA using SUMO-Ruc-soeF and Ruc-KpnI-R, and finally performing SOE PCR using these two amplification products along with primers T7-F and Ruc-Kpn-R. The SOE PCR product was then cloned into pET24b(+). pET24b(+)-HisSUMO-*ruc-*Nterm was made similarly, except that the primer RucNterm-KpnI-R substituted for Ruc-KpnI. pET24b(+)-HisSUMO-*ruc-*C29S,C32S was made by SOE PCR using primers T7-F, T7-term, Rv0991c-C29SC32S-F and Rv0991c-C29SC32S-R, with pET24b(+)-HisSUMO-*ruc* as the PCR template. pET24b(+)- HisSUMO-*ruc-*C8SC11S was made similarly to pET-24b(+)-HisSUMO-*ruc*, except that the amplification product from primers SUMO-Ruc-soeF and Ruc-KpnI-R was made using pET24b(+)-Rv0991c-C8S,C11S as a template.

Calcium chloride-competent *E. coli* DH5α was transformed with ligations. All plasmids were purified from *E. coli using* the QIAprep Spin Miniprep Kit. All plasmids made by PCR cloning were sequenced by GENEWIZ, Inc. to ensure the veracity of the cloned sequence. Plasmids were transformed into *M. tuberculosis* by electroporation as previously described (56). Single-colony transformants were isolated on 7H11 agar with antibiotic selection.

### Protein purification, antibody production, and immunoblotting

Ruc-His_6_ was produced in *E. coli* strain ER2566; His_6_-SUMO-Ruc, His_6_-SUMO-RucNterm, His_6_-SUMO- Ruc_C8S,C11S_, His_6_-SUMO-Ruc_C29S,C32S_, His_6_-SUMO-DnaK, His_6_-SUMO-DnaJ2, His_6_-SUMO-GrpE, and His_6_-Ulp1 were produced in *E. coli* strain BL21. Proteins were purified from *E. coli* by affinity chromatography using Ni-NTA agarose (Qiagen) according to the manufacturer’s instructions (Ruc-His_6_ was purified under urea denaturing conditions). Production of His_6_-SUMO-DnaK, His_6_-SUMO-DnaJ2, and His_6_-SUMO-GrpE in *E. coli* was performed as previously described (11). To make rabbit polyclonal immune serum, approximately 200 µg Ruc-His_6_ was used to immunize rabbits (Covance, Denver, PA).

*M. tuberculosis* Ruc was prepared from *E. coli* using two methods that yielded approximately equal purity; both methods also resulted in identical Ruc chaperone activity upon oxidation of the protein. In the first method, untagged Ruc was purified from strain ER2566. A 500 ml culture was grown at 37°C with shaking to an OD_600_ of 0.5, and then cooled to room temperature. 1 mM isopropyl β-D-thiogalactopyranoside (IPTG) was added, and the culture was grown further at 30°C with shaking for five hours. Bacteria were collected by centrifugation, resuspended in 25 ml of 50 mM Tris, pH 8.0, and lysed by sonication. After removing insoluble material by centrifugation, ammonium sulfate was added to the lysate to 70% w/v, and the suspension was stirred for 30 minutes at 4°C to precipitate proteins. The precipitate was collected by centrifugation, resuspended in 3 ml of 50 mM Tris, pH 8.0, and dialyzed against four liters of the same buffer at 4°C overnight to remove residual ammonium sulfate and dissolve proteins. Ruc has a predicted isoelectric point of approximately 9, and was therefore one of the few positively-charged proteins in the bacterial protein extract. Taking advantage of this fact, Ruc was purified to homogeneity by passing the protein extract over a Q Sepharose anion exchange column (GE); Ruc immediately exited the column in flow-through fractions. To ensure that the protein was fully reduced after purification, we treated Ruc with 5 mM DTT for 30 minutes at 30°C. An Amicon centrifugal filter unit (Millipore) was used to thoroughly buffer-exchange the protein into 50 mM HEPES, pH 7.5. The complete removal of DTT was confirmed by measuring the presence of DTT in the filter flowthrough using Ellman’s reagent (Thermo Scientific) according to the manufacturer’s instructions. The final protein preparation (Ruc_red_) was stored in the same buffer at −20°C.

The second method of preparing Ruc from *E. coli* used a cleavable affinity tag, and was also used to obtain Ruc_Nterm_, Ruc_C8S,C11S_, and Ruc_C29S,C32S_. His_6_-SUMO-Ruc, His_6_-SUMO-Ruc_Nterm_, His_6_-SUMO-Ruc_C8S,C11S_, and His_6_-SUMO-Ruc_C29S,C32S_ were each purified from strain BL21(DE3) in the following manner. A 500 ml culture was grown at 37°C with shaking to an OD_600_ of 0.3 to 0.4, then transferred to a 25°C shaking incubator and grown to an OD_600_ of 0.5. 1 mM IPTG was added, and the culture were further grown for 5 hours. Purified protein was prepared from bacteria using Ni-NTA resin, and then exchanged into a SUMO cleavage buffer of 50 mM Tris, 150 mM NaCl, 2 mM dithiothrietol (DTT), pH 8.0. A 1:100 volume of purified His_6_-Ulp1 (SUMO protease) was added, and the reaction was incubated for 30 minutes at 30°C. His_6_-Ulp1 and His_6_-SUMO were then removed by incubating the mixture with Ni-NTA resin and saving the supernatant fraction; clearance of tagged protein was performed twice to yield pure, native Ruc and truncation or substitution variants. Proteins were buffer-exchanged into TBS buffer (50 mM Tris, 150 mM NaCl, pH 8.0) using centrifugal filters, and the absence of residual DTT was confirmed using Ellman’s reagent. The final protein preparations (Ruc_red_, Ruc_Nterm-red_, Ruc_C8S,C11S-red_, and Ruc_C29S,C32S-red_) were stored in TBS at −20°C.

*M. tuberculosis* DnaK, DnaJ2, and GrpE were prepared by removing the affinity tags from His_6_-SUMO-DnaK, His_6_-SUMO-DnaJ2, and His_6_-SUMO-GrpE in the same manner as described for His_6_-SUMO-Ruc, except that the native proteins were buffer-exchanged into 50 mM Tris, 150 mM KCl, 20 mM MgCl2, 2 mM DTT, pH 7.5 before storage at −20°C.

Separation of proteins in *in vitro* assays and in *M. tuberculosis* lysates was performed using 15% sodium dodecyl sulfate-polyacrylamide (SDS-PAGE) gels, with the following exceptions: for Figures 1B and 1C, anykD Mini-PROTEAN TGX precast protein gels (Bio-Rad) were used. Bio-Safe Coomassie Stain (Bio-Rad) was used to stain gels. For preparing samples for SDS-PAGE gels in Figures 3A and 4B, purified proteins were mixed with 4 × non-reducing SDS buffer (250 mM Tris pH 6.8, 2% SDS, 40% glycerol, 1% bromophenol blue) to a 1 × final concentration, and samples were boiled for five minutes. For lane 3 in Figure 3A, DTT was added to the sample to a 2 mM final concentration prior to boiling. For immunoblots, proteins were transferred from protein gels to nitrocellulose membranes (GE Amersham), and analyzed by immunoblotting as indicated. Due to Ruc’s high isoelectric point, Ruc was transferred to membranes using 100 mM N-cyclohexyl-3-aminopropanesulfonic acid (CAPS) buffer in 10% methanol. In Figures 1B and 1C, Ruc and PrcB immunoblots were from the same membrane. For detecting *M. tuberculosis* DnaK we used a monoclonal antibody from BEI Resources (NR-13609) at a concentration of 1:1000 in 3% BSA in 25 mM Tris-Cl/125 mM NaCl/0.05% Tween 20 buffer (TBST). Polyclonal antibodies against PrcB (23) and Ruc were used similarly. Secondary antibodies HRP-conjugated goat anti-rabbit IgG F(ab’)2 and HRP-conjugated anti-mouse IgG(H+L) were purchased from Thermo Fisher Scientific. All antibodies were made in TBST with 3% BSA. Immunoblots were developed using SuperSignal West Pico PLUS chemiluminescent substrate (Thermo Fisher Scientific) and imaged using a Bio-Rad ChemiDoc system.

### Preparation of *M. tuberculosis* extracts for immunoblotting

*M. tuberculosis* cultures were grown to an OD_580_ of 0.3. Equal amounts of bacteria were harvested by centrifugation, resuspended in TBS, and transferred to a tube containing 250 µl of 0.1 mm zirconia beads (BioSpec Products). Bacteria were lysed using a mechanical bead-beater (BioSpec Products). Whole lysates were mixed with 4 × reducing SDS sample buffer (250 mM Tris pH 6.8, 2% SDS, 20% 2-mercaptoethanol, 40% glycerol, 1% bromophenol blue) to a 1 × final concentration, and samples were boiled for 5 minutes. For preparing lysates from *M. tuberculosis* grown in 7H9, which contains BSA, an additional wash step with PBS-T was done prior to resuspension of bacteria in lysis buffer.

### Mouse infections

Six to eight-week old female C57BL6/J mice (Jackson Laboratories) were each infected with ∼200-400 *M. tuberculosis* bacilli by the aerosol infection route. Bacterial burden in organs was determined as previously described (57); briefly, at the time points indicated in the text, lung pairs and spleens were harvested from 3 to 5 mice, and were each homogenized and plated on 7H11 agar to determine CFU per organ. All procedures were performed with the approval of the New York University Institutional Animal Care and Use Committee.

### Preparation of Ruc_ox_ and Ruc_Nterm-ox_

8.8 M H_2_O_2_ stock solution, stored at −20°C, was diluted to 200 mM in deionized water just prior to the oxidation reaction. 100 µM Ruc_red_ or Ruc_Nterm-red_ were incubated at 37°C; for 3 min, after which 50 µM of copper chloride was added, followed by 2 mM H_2_O_2_. The reaction was incubated for 10 minutes; H_2_O_2_ and copper chloride were removed using a Zeba Spin 7K MWCO desalting centrifuge column (Fisher Scientific) pre-equilibrated with the original Ruc_red_ storage buffer, according to the manufacturer’s instructions. Treatment of Ruc with 2 mM DEANO (Sigma-Aldrich) or NaOCl (Figure S1B) was performed identically, except that Ruc was treated for 20 minutes (NaOCl) or two hours (DEANO) as described for Hsp33 (58).

### Spectrophotometric detection of zinc in Ruc preparations

Quantification of the zinc coordinated by Ruc cysteines was performed using 4-(2-pyridylazo)resorcinol (PAR) using a previously established method (32, 59), except that zinc-cysteine complexes were disrupted using N-ethylmaleimide (NEM) (Pierce) according to the manufacturer’s instructions. 25 µM Ruc was mixed with 100 µM PAR either in the absence or presence of 2 mM NEM. Reactions were incubated at room temperature for 1 hour. Zn(PAR)_2_ complexes were detected by A_500_ using a NanoDrop spectrophotometer. To obtain a precise concentration of zinc in protein preparations, serial dilutions of zinc sulfate prepared in matched buffers (with or without NEM) were used to prepare standard curves. Three technical replicates were performed for each condition.

### Metal analysis of Ruc preparations using inductively-coupled plasma mass spectrometry (ICP-MS)

Elemental quantification on purified Ruc with and without NEM and a buffer control was performed using an Agilent 7700 inductively coupled plasma mass spectrometer (Agilent, Santa Clara, CA) attached to a Teledyne CETAC Technologies ASX-560 autosampler (Teledyne CETAC Technologies, Omaha, NE). The following settings were fixed for the analysis Cell Entrance = −40 V, Cell Exit = −60 V, Plate Bias = −60 V, OctP Bias = −18 V, and collision cell Helium Flow = 4.5 mL/min. Optimal voltages for Extract 2, Omega Bias, Omega Lens, OctP RF, and Deflect were determined empirically before each sample set was analyzed. Element calibration curves were generated using ARISTAR ICP Standard Mix (VWR, Radnor, PA). Samples were introduced by peristaltic pump with 0.5 mm internal diameter tubing through a MicroMist borosilicate glass nebulizer (Agilent). Samples were initially up taken at 0.5 rps for 30 s followed by 30 s at 0.1 rps to stabilize the signal. Samples were analyzed in Spectrum mode at 0.1 rps collecting three points across each peak and performing three replicates of 100 sweeps for each element analyzed. Sampling probe and tubing were rinsed for 20 s at 0.5 rps with 2 % nitric acid between every sample. Data were acquired and analyzed using the Agilent Mass Hunter Workstation Software version A.01.02.

### Circular dichroism (CD) spectrophotometry

CD measurements were performed using a Jasco J-1500 CD spectrophotometer as per the manufacturer’s instructions. To prepare samples, Ruc was buffer-exchanged into 20 mM KH_2_PO_4_, pH 7.5, diluted to 20 µM, and transferred to a quartz cuvette. The spectrophotometer parameters were set as follows: CD scale 200 mdeg/1.0 dOD, integration time 1 second, bandwidth 1 nm. The voltage was monitored simultaneously and remained below 700 V.

### Luciferase aggregation and refolding assays

The chaperone activity of Ruc was measured by testing its ability to limit the aggregation of heat-denatured luciferase, a previously established method (15, 37). For experiments in Figures 4A, 4C, and 4E, aggregation was determined by measuring absorbance at 350 nm (A_350_) (60, 61). Firefly luciferase (Promega) was diluted to 2 µM in TBS and mixed in a microcentrifuge tube with concentrations of Ruc indicated in the text and figures, or TBS alone, to a final volume of 20 µl. The tube was then incubated in a 45°C heat block. Immediately before incubation and at the indicated times, a 2 µl volume was removed and A_350_ was measured using a NanoDrop spectrophotometer. Aggregation assays were performed using three technical replicates per condition. For the experiment in Figure S1A, luciferase aggregation was measured using light scattering as previously described (15).

Refolding of heat-denatured luciferase by *M. tuberculosis* DnaKJE was performed using a protocol adapted from (11). *M. tuberculosis* encodes two DnaJ homologs, DnaJ1 and DnaJ2, which both promote DnaK-mediated protein folding *in vitro* (11); DnaJ2 was used in this study because we found that DnaJ1 exhibited poor solubility when purified from *E. coli*. For the denaturation step, luciferase was diluted to 0.1 µM in TBS, either alone or mixed with 4 µM Ruc_red_ or Ruc_ox_, in glass vials. Denaturation was performed by placing vials in a 45°C water bath for 20 minutes. For the refolding step, we used glass-coated 96-well plates in which 5 µl of denaturation reaction was mixed with 15 µl of refolding reaction buffer [50 mM Tris pH 7.5, 150 mM KCl, 20 mM MgCl_2_, 2 mM DTT, 1 mg/ml BSA (Sigma-Aldrich), 2 mM Mg_2+_-ATP] or refolding reaction buffer containing 4 µM DnaK, 2 µM DnaJ2, and 2 µM GrpE. Plates were incubated at 25°C. Luciferase activity was measured immediately upon initiating the refolding step (0 minutes) and at all other time points indicated in Figure 5A by transferring 2 µl of each reaction into a white opaque 96-well plate and adding 100 µl of luciferase assay mix (100 mM KH_2_PO_4_ pH 7.5, 25 mM glycyl glycine, 0.2 mM EDTA, 2 mM Mg_2+_-ATP, 0.5 mg/ml BSA, 70 µM luciferine). Luminescence was measured using a plate reader (PerkinElmer EnVision). For calculating the percent of native luciferase activity for all reactions at each time point, a control reaction was included in which 0.1 µM luciferase was diluted into TBS in a glass vial but kept at 4°C, rather than heated, during the denaturation step. This control reaction was mixed with refolding reaction buffer as described above, and luminescence was measured at each time point in Figure 5A to determine 100% native luciferase activity.

### Tandem affinity purification of *M. tuberculosis* proteins

Purifications of TAP-tagged proteins from *M. tuberculosis* were performed under low-salt conditions as described (62). The following changes were made to the protocol: 100 µl of packed Ni-NTA beads and 100 µl of M2 anti-FLAG affinity gel were used; 100 µl of 100 µM 3X FLAG peptide was used for the final elution. For capturing Ruc_TAP_ interactions with other *M. tuberculosis* proteins, *M. tuberculosis* lysates were incubated at 45°C for 10 minutes either in the absence (Figure 5C, lane 1) or presence (Figure 5C, lane 2) of 2 mM H_2_O_2_ and 50 µM CuCl_2_; purifications were subsequently performed as described above. Samples were boiled in 4 × reducing SDS sample buffer prior to running SDS-PAGE gels.

### Protein mass spectrometry

To identify of the ∼70 kDa protein pulled down by Ruc_TAP_ (Figure 5B), the band was excised from a SDS-PAGE gel and processed as previously described (63). To determine the identity of proteins enriched in Ruc_TAP_ purifications under oxidizing conditions (Figure 5C; Table S2), Ruc_TAP_ purifications under native or oxidizing conditions were performed in three biological replicates. Affinity purified samples were reduced, alkylated, digested with trypsin and desalted as previously described (23, 63). The peptide eluates in all cases were concentrated in the SpeedVac and stored at - 80°C. Aliquots of each sample were loaded onto a trap column (Acclaim® PepMap 100 pre-column, 75 μm × 2 cm, C18, 3 μm, 100 Å, Thermo Scientific) connected to an analytical column (EASY-Spray column, 50 m × 75 μm ID, PepMap RSLC C18, 2 μm, 100 Å, Thermo Scientific) using the autosampler of an Easy nLC 1000 (Thermo Scientific) with solvent A consisting of 2% acetonitrile in 0.5% acetic acid and solvent B consisting of 80% acetonitrile in 0.5% acetic acid. The peptide mixture was gradient eluted into the Orbitrap QExactive HF-X mass spectrometer (Thermo Scientific) using the following gradient: 5%-35% solvent B in 120 min, 35% −45% solvent B in 10 min, followed by 45%-100% solvent B in 20 min. The full scan was acquired with a resolution of 45,000 (@ m/z 200), a target value of 3e6 and a maximum ion time of 45 ms. Following each full MS scan, twenty data-dependent MS/MS spectra were acquired. The MS/MS spectra were collected with a resolution of 15,000 an AGC target of 1e5, maximum ion time of 120ms, one microscan, 2m/z isolation window, fixed first mass of 150 m/z, dynamic exclusion of 30 sec, and Normalized Collision Energy (NCE) of 27. All acquired MS2 spectra were searched against a UniProt *M. tuberculosis* H37Rv database including common contaminant proteins using Sequest HT within Proteome Discoverer 1.4 (Thermo Fisher Scientific). The search parameters were as follows: precursor mass tolerance ±10 ppm, fragment mass tolerance ±0.02 Da, digestion parameters trypsin allowing two missed cleavages, fixed modification of carbamidomethyl on cysteine, variable modification of oxidation on methionine, and variable modification of deamidation on glutamine and asparagine and a 1% peptide and protein FDR searched against a decoy database. The results were filtered to only include proteins identified by at least two unique peptides. Fold change analysis was performed for the Ruc_TAP_ purifications using the ratios of PSMs in oxidized to the PSMs in the native affinity purified samples using the SAINT algorithm (64). SAINT scores were used to calculate the false discovery rate (FDR); proteins whose SAINT score yielded an FDR of 5% or lower were considered statistically significant and are highlighted in Table S2.

### Measurement of *M. tuberculosis* susceptibility to oxidants

*M. tuberculosis* was grown in 7H9c to an OD_580_ of 0.5 at 37°C, centrifuged and resuspended in fresh 7H9c, and spun at 500 × g to remove clumps of bacteria. Supernatants were then diluted to an OD_580_ of 0.025, transferred to 96-well plates, and incubated at 45°C for four hours to induce Ruc production. Afterwards, *M. tuberculosis* strains were subjected to oxidizing reagents and inoculated onto 7H11 agar as described in the legend for Fig. S3.

### Computational analyses

Iterative sequence profile searches were performed to recover Ruc sequence homologs using the PSI-BLAST program (65). Searches were either run against the non-redundant (nr) protein database of the National Center for Biotechnology Information (NCBI), or a custom database of 7423 complete prokaryotic genomes extracted from the NCBI Refseq database (66). The latter was used for phyletic profile analyses (Table 2). Contextual information from prokaryotic gene neighborhoods was retrieved using a Perl script that extracts the upstream and downstream genes of a query gene from a GenBank genome file. This was followed by clustering of proteins using the BLASTCLUST program (ftp://ftp.ncbi.nih.gov/blast/documents/blastclust.html) to identify conserved gene-neighborhoods. Analysis and visualization of phyletic patterns was performed using the R language.

**Table 2.** Occurance of *ruc* in bacterial and archaeal phyla.

| **Bacterial phylum (number of genomes)** | **% genomes with *ruc*** |
| --- | --- |
| gammaproteobacteria (923) | 26.65 |
| betaproteobacteria (333) | 82.88 |
| zetaproteobacteria (5) | 80 |
| alphaproteobacteria (469) | 17.27 |
| deltaproteobacteria (86) | 86.05 |
| proteobacteria (43) | 74.42 |
| thermodesulfobacteria (4) | 100 |
| spirochaetes (70) | 48.57 |
| defferibacteres (4) | 100 |
| chrysiogenetes (1) | 100 |
| nitrospirae (73) | 86.3 |
| acidobacteria (9) | 100 |
| elusimicrobia (3) | 33.33 |
| verrucomicrobia (74) | 60.81 |
| chlamydiae (43) | 6.98 |
| planctomycetes (18) | 100 |
| bacteroidetes (263) | 1.9 |
| chlorobi (263) | 5.7 |
| fibrobacteres (2) | 0 |
| gemmatimonadetes (3) | 100 |
| ignavibacteriae (2) | 100 |
| aquificae (15) | 20 |
| dictyoglomi (2) | 100 |
| thermotogae (29) | 100 |
| deinococcus-thermus (26) | 100 |
| synergistetes (5) | 0 |
| fusobacteria (16) | 0 |
| thermobaculum (1) | 100 |
| actinobacteria (497) | 73.84 |
| chloroflexi (22) | 100 |
| armatimonadetes (15) | 73.33 |
| tenericutes (124) | 0 |
| firmicutes (772) | 8.94 |
| cyanobacteria (127) | 13.39 |
| calditrichaeota (2) | 100 |
| unclassified bacteria (2565) | 11.89 |
| **Archaeal phylum (number of genomes)** | **% genomes with *Ruc*** |
| crenarchaeota (61) | 1.64 |
| euryarchaeota (187) | 6.42 |
| archaea (31) | 3.23 |

## Acknowledgments

We thank C. Nathan and K. Burns-Huang for providing strains and technical advice. This work was supported by NIH grants AI088075 and AI144851 awarded to K.H.D. S.H.B was supported by NIH grant T32 AT007180. S.H.B. also received support from the Jan T. Vilcek Endowed Fellowship Fund. E.P.S. received support from AI150701 and AI069233. K.U. and U.J. thank the DFG Priority Program SPP 1710 (Schw823/3-2). L.M.I. and L.A. are supported by the intramural funds of the National Library of Medicine, NIH. The mass spectrometry experiments were in part supported by the NYU Grossman School of Medicine.

## Author contributions

S.H.B., K.U., A.D., B.U., W.B., E.P.S., L.I., L. A., U.J. and K.H.D. designed research; S.H.B., K.U., A.D., W.B., L.M.I., and L.A. performed research; S.H.B, K.H.D., and L.A. wrote the manuscript.

---

**Fig. S1.**
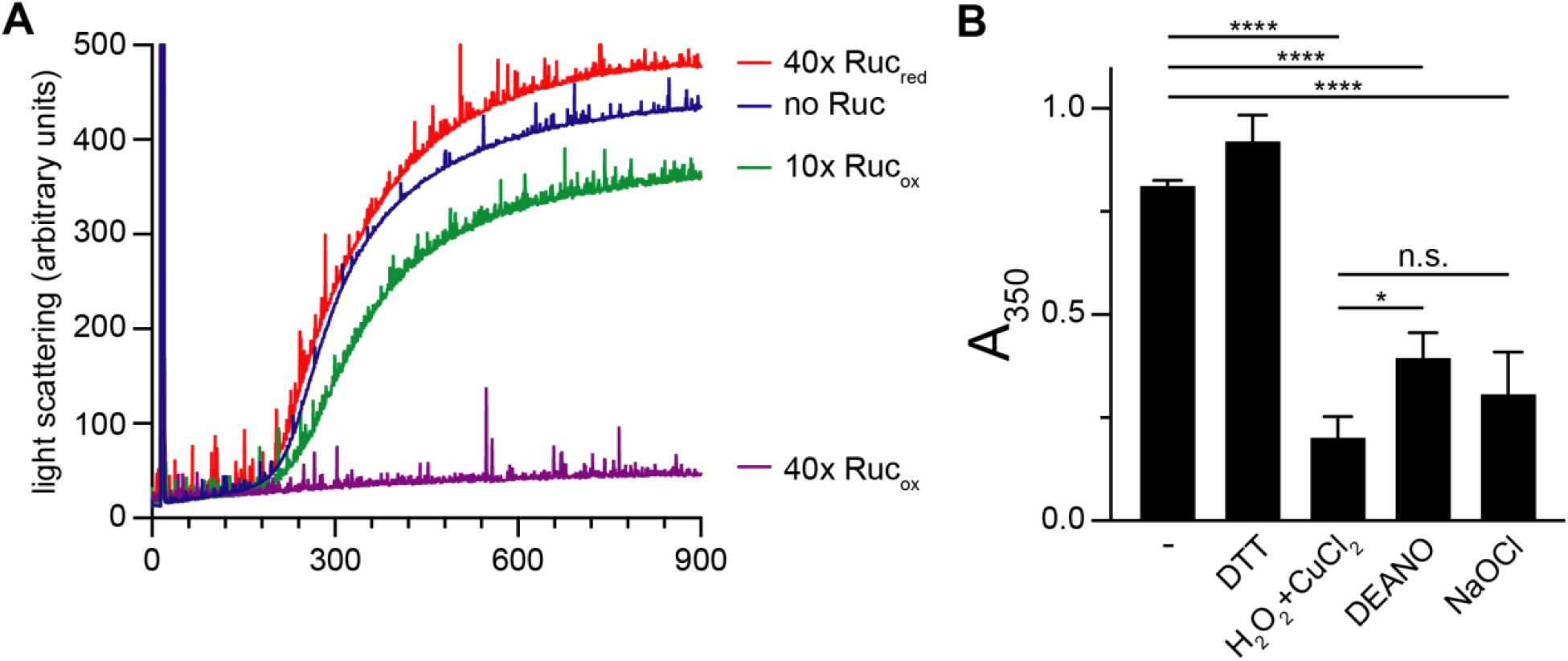
Ruc_ox_ inhibits protein aggregation and is activated by multiple oxidants. **(A)** Measurement of denatured luciferase aggregation using light scattering spectrophotometry. Luciferase was incubated alone, in the presence of a 40-fold (40x) or 10-fold (10x) molar excess of Ruc_ox_, or a 40-fold molar excess of Ruc_red_. **(B)** Measurement of denatured luciferase aggregation using absorbance at 350 nm (A_350_). Luciferase was incubated at 45°C for 5 minutes either alone (-) or in the presence of Ruc pre-treated with 2 mM DTT, 2 mM H_2_O_2_ with 0.5 mM CuCl_2_, 2 mM diethylamine NONOate (DEANO, a nitric oxide donor), or 2 mM sodium hypochlorite (NaOCl). Data in (A) is representative of two independent experiments; (B) was performed using three replicates per condition. Statistical significance was determined using one-way ANOVA; ****, *p* < 0.0001; *, *p* < 0.05; n.s., not statistically significant (*p* > 0.05).

**Fig. S2.**
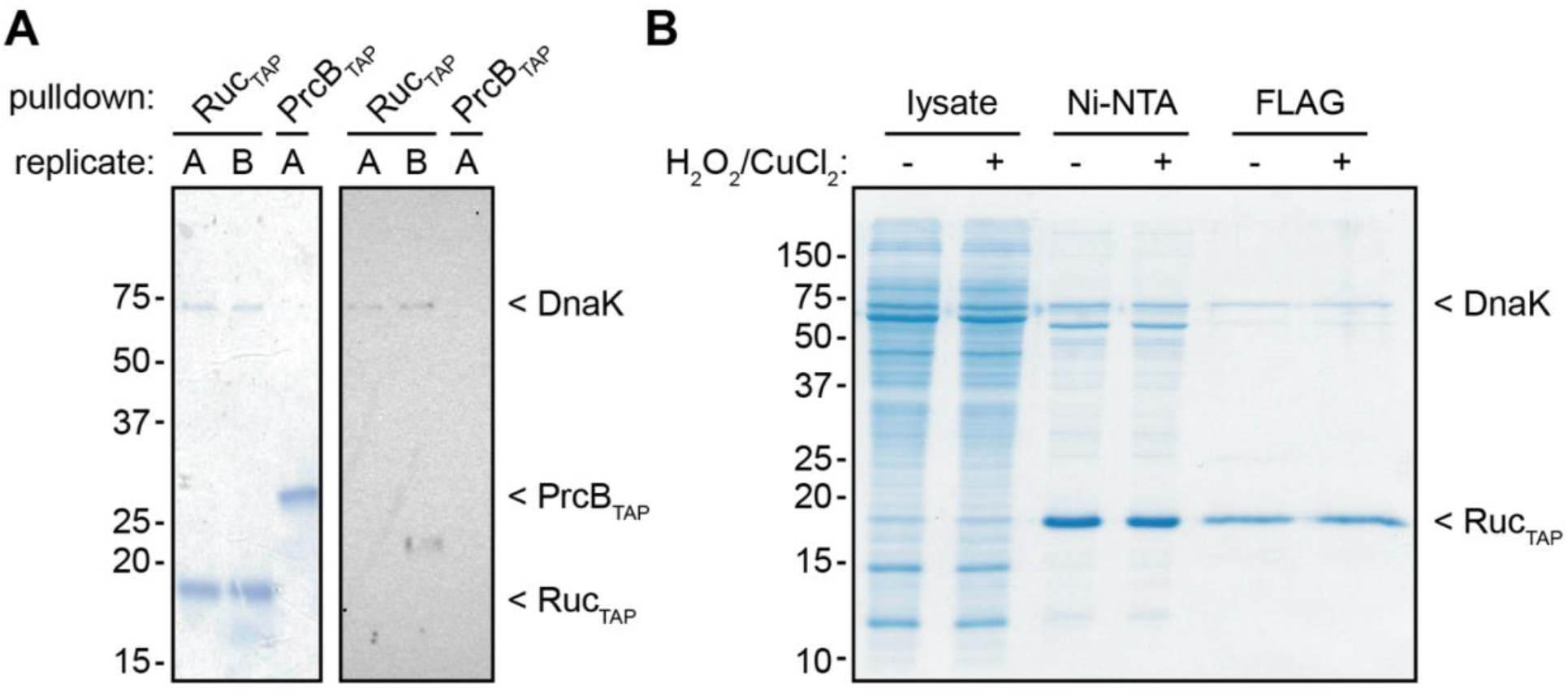
Ruc_TAP_ co-purifies with DnaK, and subjecting *M. tuberculosis* to oxidation does not affect Ruc_TAP_ purification. **(A)** Ruc_TAP_ or PrcB_TAP_ (a control, TAP-tagged protein) purifications from *M. tuberculosis* were separated on an SDS-PAGE gel and either stained with Coomassie brilliant blue (left) or analyzed by immunoblotting with a monoclonal antibody against *M. tuberculosis* DnaK (right). **(B)** Ruc_TAP_ was purified in two steps (Ni-NTA resin followed by α-FLAG affinity gel) from *M. tuberculosis* cultures (strain MHD1541) that were either untreated or treated with a sublethal concentration of oxidants (0.625 mM H_2_O_2_ with 16 nmol CuCl_2_) for 30 minutes at 45°C.

**Figure S3.**
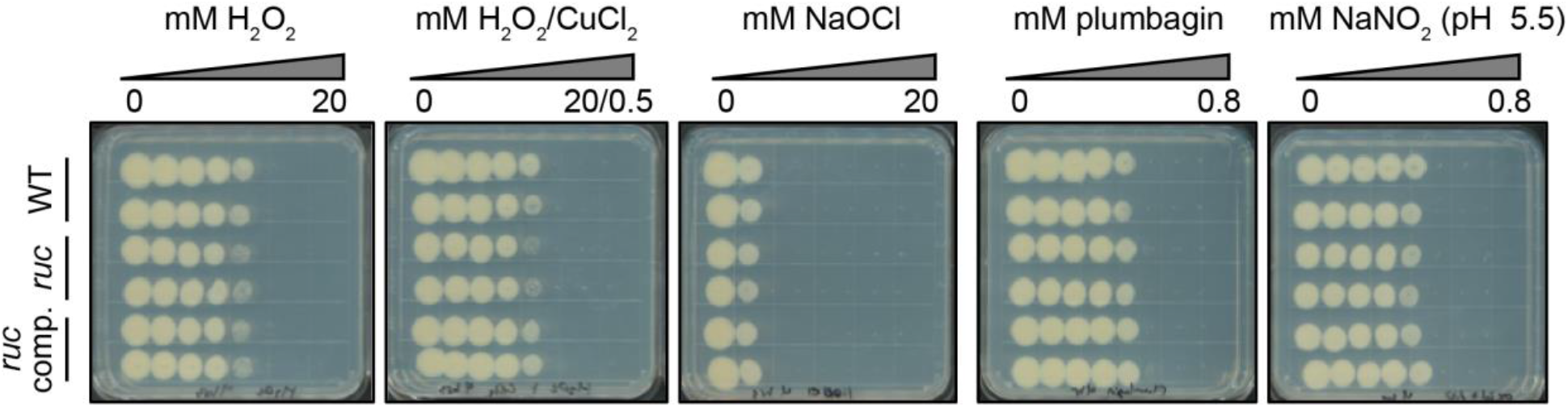
WT and *ruc* mutant strains are equally sensitive to oxidants. The approximate lethal dose of the indicated oxidizing agents (above) on *M. tuberculosis* strains (left; MHD1385, MHD1393, and MHD1394 are represented). A series of two-fold serial dilutions of each oxidant was added to *M. tuberculosis* at the final concentrations shown; cultures were incubated along with oxidants at 45°C for four hours. Afterwards, cultures were spot-plated onto solid agar media, and plates were incubated at 37°C for two weeks to assess surviving bacteria.

**Table S1.** ICP-MS analysis of Ruc metal binding.

Excel spreadsheet in separate document

**Table S2.** *M. tuberculosis* proteins that co-purified with Ruc_TAP_.

Excel spreadsheet in separate document

